# 2’, 3’, 4’-trihydroxychalcone is an Estrogen Receptor Ligand Which Modulates the Activity of 17β-estradiol

**DOI:** 10.1101/607275

**Authors:** Candice B. Herber, Jeanne G Quirit, Gary Firestone, Charles Krois

## Abstract

Menopausal hormone therapy (MHT) reduces the risk of osteoporosis, fractures, obesity and diabetes, but long-term MHT increases risk of other diseases. Safe long-term MHT that exploits its benefits and abrogates its adverse effects requires a new approach. Here we demonstrate that 2’, 3’, 4’-trihydroxychalcone (CC7) acts as an estrogen receptor alpha (ERα) ligand that may improve the safety profile of MHT. CC7 reprograms the actions of estradiol (E2) to regulate unique genes in bone-derived U2OS cells, with 824/1358 genes not regulated by E2. The proliferative action of E2 on human MCF-7 breast cancer cells and mouse uterus is blocked when combined with CC7. Thermostability and molecular dynamics simulation studies suggest that CC7 binds concurrently with E2 in the ERα ligand binding pocket to produce a unique coliganded conformation to modulate ERα. Compounds such as CC7 that act as coligands represent a new class of ERα reprograming drugs that potentially can be combined with existing estrogens to produce a safer MHT regimen for long-term therapy.

## INTRODUCTION

Reduced estrogen production during menopause leads to short-term symptoms of hot flashes and vaginal atrophy, and long-term increases in osteoporosis, bone fractures, weight gain and type 2 diabetes^6^. Menopausal hormone therapy (MHT) uses estrogen treatment to prevent both short and long-term symptoms^6^.The original MHT regimen contained only estrogens, which act through a one-ligand, one-receptor class mechanistic model: a single estrogen molecule binds to the ligand binding domain (LBD) of ERα or ERβ^1^. Unfortunately, estrogens in MHT promote uterine cell proliferation by activating ERα, increasing the risk of endometrial cancers^2^. To overcome the proliferative effects on the endometrium, postmenopausal women are prescribed a MHT combination regimen of estrogen and a progestin^3^. This combination acts through a two-ligand, two-receptor mechanism; progestin binds to the progesterone receptor, which inhibits the proliferative action of estrogen-bound ERα in the uterus, thus eliminating the risk of endometrial cancer^3,4^. However, long-term combination MHT increases the incidence of breast cancer, cardiovascular disease and Alzheimer’s in postmenopausal women and is contraindicated because of these risks^45^.

To target the long-term beneficial indications of MHT, it will be necessary to develop safer regimens. Currently, there are two strategies. First, estrogens have been combined with the selective estrogen receptor modulator (SERM), bazedoxifene, which prevents estrogen binding to ERα, thereby blocking the proliferative effects on the uterus^7^. This combination is approved only for short-term treatment of hot flashes. Second, ERβ-selective agonists have been developed^8^, which are promising due to the antiproliferative action of ERβ in breast cancer cells^9^. However, because ERα is the major ER in bone and adipose tissue, these drugs may not be effective for osteoporosis, weight gain or diabetes^10–12^. In addition, ERβ selective agonists are not approved for clinical use. Since no drugs are approved for long-term MHT therapy, we screened compounds that could bind concurrently with E2 to ERα, acting as coligands at a secondary site within the LBD creating a two-ligand, one-receptor mechanistic approach. We identified 2’, 3’, 4’-THC (referred to as CC7) as an ER coligand that modulates the activity of E2 bound to ERα on gene expression and cell proliferation. CC7 could potentially be combined with existing estrogens to make MHT safe for continuous, long-term administration to prevent chronic diseases associated with menopause.

## RESULTS

### CC7 acts as an ER coagonist in U2OS cells

Previously, we isolated the flavanone liquiritigenin from the plant Glycyrrhizae uralensis Fisch and showed that it is an ERβ-selective agonist^13^. To identify compounds that alter the activity of E2 by acting as a coligand, we leveraged a unique screening method (Fig 1A) whereby we screened compounds structurally similar to liquiritigenin in transfection assays for their ability to synergize with E2 (Fig 1B). CC7 (Fig. 1C) was selected for further studies based on our initial screening. In U2OS osteosarcoma cells transfected with ERα or ERβ, CC7 was inactive on an ERE upstream of the luciferase gene, whereas it produced a synergistic increase in luciferase activity with E2 in the presence of both ERs (Fig. 1D). We focused on the effects of CC7 on ERα activity, because it is the main mediator of deleterious effects with long-term MHT. To explore synergy between CC7 and E2 on endogenous ERα target genes, we measured their effects on keratin 19 (KRT19)^14^. Both 1 and 5 μM CC7 activated KRT19 only in the presence of E2 (Fig. 1E). Since the synergy trended greater with 5 μM we used this concentration in subsequent studies. Using stably transfected U2OS-ERα cells, we performed competition binding studies to determine if CC7 competes with [3H]-E2. Surprisingly, 5 and 10 μM CC7 produced an approximate 2-fold increase in [3H]-E2 binding (Fig. 1F and S1A). Competition occurred only at 100 μM CC7 (S1A), a 100-fold higher concentra-tion than required for synergy (Fig. 1E).

**Figure 1.**
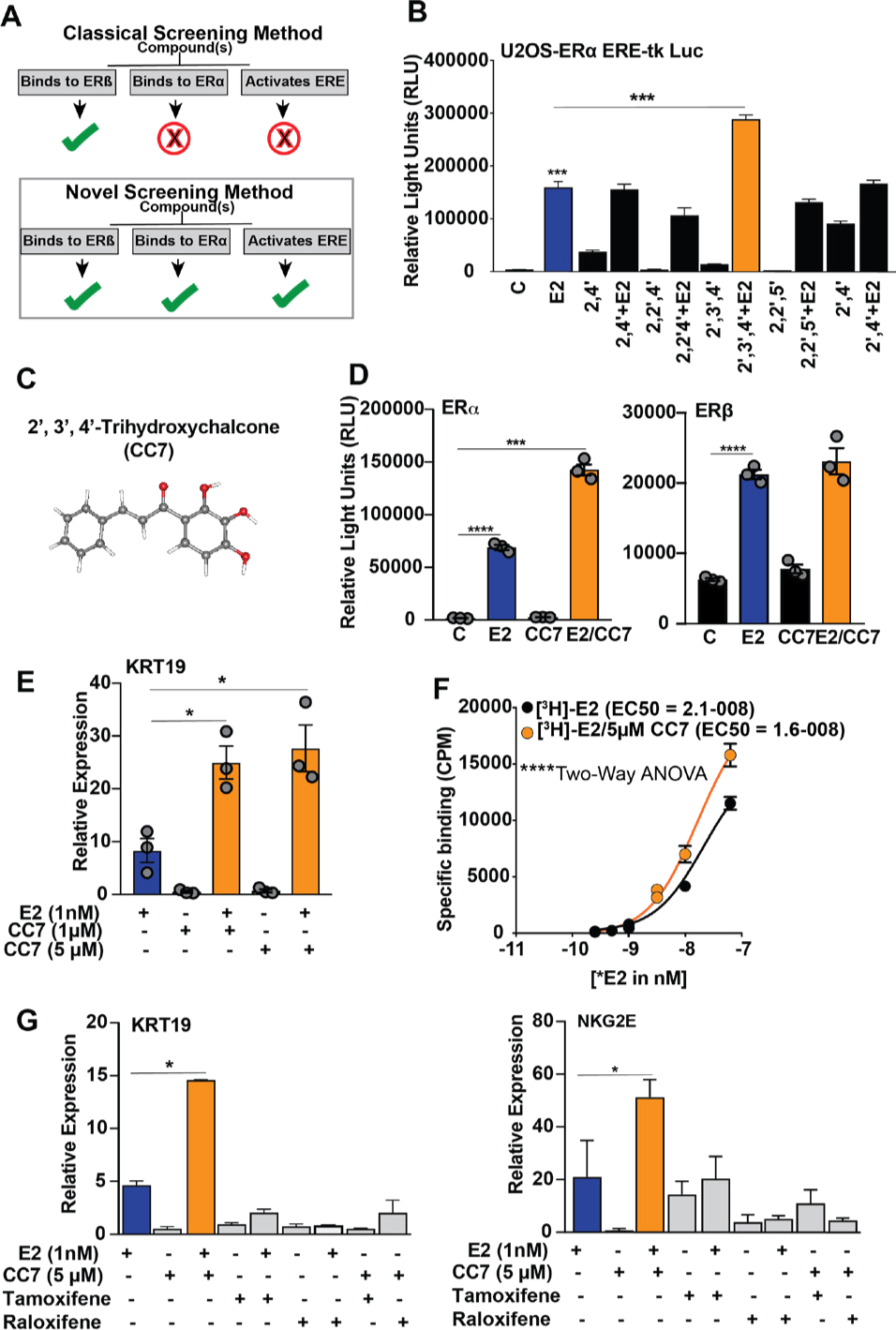
CC7 synergizes the estradiol (E2) induced transcriptional activities mediated by ERs. (**A**) Schematic of novel screening method (**B**) Luciferase activities of U2OS cells cotransfected with ERE tk-luciferase and ERα and treated with chalcone compounds for 24h. (**C**) Chemical structure of CC7 adapted from PubChem (**D**) Luciferase activities of U2OS cells cotransfected with ERE tk-luciferase and ERα (left panel) or ERβ (right panel). Cells were treated with 1 nM of E2 and 5 μM of 2’, 3’, 4’-THC alone or in combination for 24 hours. (**E**) KRT19 mRNA levels in U2OS-ERα cells treated with 1 nM E2 in the absence or presence of 1 or 5 μM CC7 for 24 hrs. (**F**) Competitive binding curve in U2OS-ERα cells treated with 5 μM CC7 and increasing doses of E2 for 1 hr at 37°C. Total [3H]-E2 binding was measured with a scintillation counter. (**G**) KRT-19 and NKG2E gene expression in U2OS-ERα cells treated with 1 nM E2, 5 μM CC7 in the presence and absence of 1 μM of tamoxifen or raloxifene for 24 hours determined by RT-qPCR. The data shown are mean ± SEM of technical triplicates. The asterisks indicate the significant difference between E2 alone and E2 in combinations. Statistical significances of all data in the figure were analyzed with one-way ANOVA followed by Tukey’s multiple comparisons post hoc test.

### CC7 behaves differently than SERMs in U2OS-ERα cells

To compare the activity of CC7 to SERMs on endogenous gene regulation, we examined their effects on the expression of ERα targets KRT19^14^ and NKG2E^15^. E2 induced KRT19 mRNA expression in U2OS-ERα cells compared to the vehicle control, whereas no effect was observed with CC7 or the SERMS tamoxifen or raloxifene (Fig. 1G). The CC7/E2 combination produced a synergistic activation of KRT19, whereas tamoxifen and raloxifene blocked E2-induced expression (Fig. 1G). E2, tamoxifen and raloxifene activated NKG2E mRNA expression, whereas no effect was observed with CC7 (Fig. 1G). A synergistic activation of the NKG2E gene occurred with the CC7/E2 combination, whereas raloxifene antagonized the E2 effect. These data demonstrate that CC7 does not behave like classical SERMs in U2OS cells.

To determine if CC7 alters E2 regulation of other genes, we performed microarray analysis in U2OS-ERα cells. At 10 nM E2 alone regulated 756 genes, whereas CC7 alone regulated only 31 genes, of which 14 genes were regulated by E2 and 25 genes were regulated by the CC7/E2 combination (Fig 2A and Supplementary Table S1). The CC7/E2 combination regulated 1,358 genes, and of these genes 534 were also regulated by E2 (Supplementary Table S1). 824 genes were termed unique genes (Supplementary Table S1) because they were only regulated by the CC7/E2 combination (Fig. 2A). To verify the microarray results, we examined the activation of some of the unique genes including FGR, KCNK6 and K6IRS3 (Fig. 2B). KCNK6 and K6iRS3 were not appreciably regulated by E2, whereas FGR was activated at 10 nM E2. Similar to the microarray data, the CC7/E2 combination produced a large, synergistic activation of the KCNK6, K6IRS3 and FGR genes at 10 nM E2. The ER antagonist ICI 182,780 blocked the induction of unique genes (Supplementary Fig. S1B) and NKG2E (Supplementary Fig. S1C) genes by the CC7/E2 combination, demonstrating that synergy requires ERα. Since CC7 increased E2 binding, we assayed recruitment of ERα the KRT19 gene. By ChIP, the CC7/E2 combination increased recruitment of ERα versus E2 alone (Fig. 2C).

**Figure 2.**
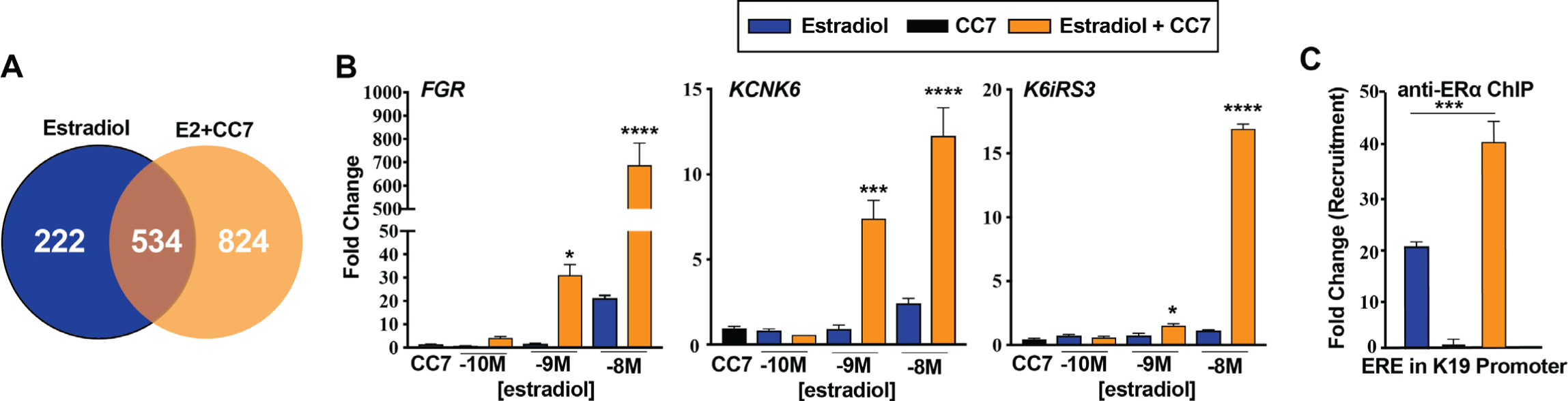
CC7 behaves as a unique coagonist of ERα on gene expression in U2OS-ERα cells. (**A**) Venn-diagram showing the total number of genes uniquely regulated by 10 nM E2 alone or in combination with 5 μM CC7. Expression was assessed by up regulation or down regulation of 3-fold or more and a p value ≤ 0.05 determined by Student t-test. Gene expression of uniquely regulated genes of (**B**) FGR KCNK6 and K6IRS3 in U2OS-ERα cells treated with increasing doses of E2 alone or in combination with 5 μM CC7 for 24 hours determined by RT-qPCR. The asterisks indicate the significance between various amounts of E2 alone and corresponding combination, which was analyzed with Two-way ANOVA followed by Sidak’s multiple comparisons post hoc test. (**C**) ERα recruitment to K19 ERE ERα binding site examined by ChIP assay after the cells were treated with E2 alone and in combination with CC7 for 2 hrs. The data shown are mean ± SEM done in technical triplicates. The statistical significance was determined by one-way ANOVA followed by Tukey’s multiple comparisons post hoc test.

### CC7 is a coligand with E2 that binds to the ERα LBD by modeling studies

During our initial screen, we found a structurally similar chalcone, 2, 2’, 4’-THC (referred to as CC6) (Fig 3A) did not produce the synergistic activation observed with CC7 (data not shown). We hypothesized that both CC6 and CC7 bind to the same site on ERα, with similar affinities. Thus, CC6 might be able to block the synergy by competing with CC7 at its binding site. If E2 and CC7 are truly co-ligands, binding at different sites, CC6 will not compete with E2, and E2 activation should be preserved. As predicted by our model, increasing concentrations of CC6 blocked the synergistic effect of CC7 on KRT-19 gene expression (Fig. 3B), but had no effect on the activation by E2.

**Figure 3.**
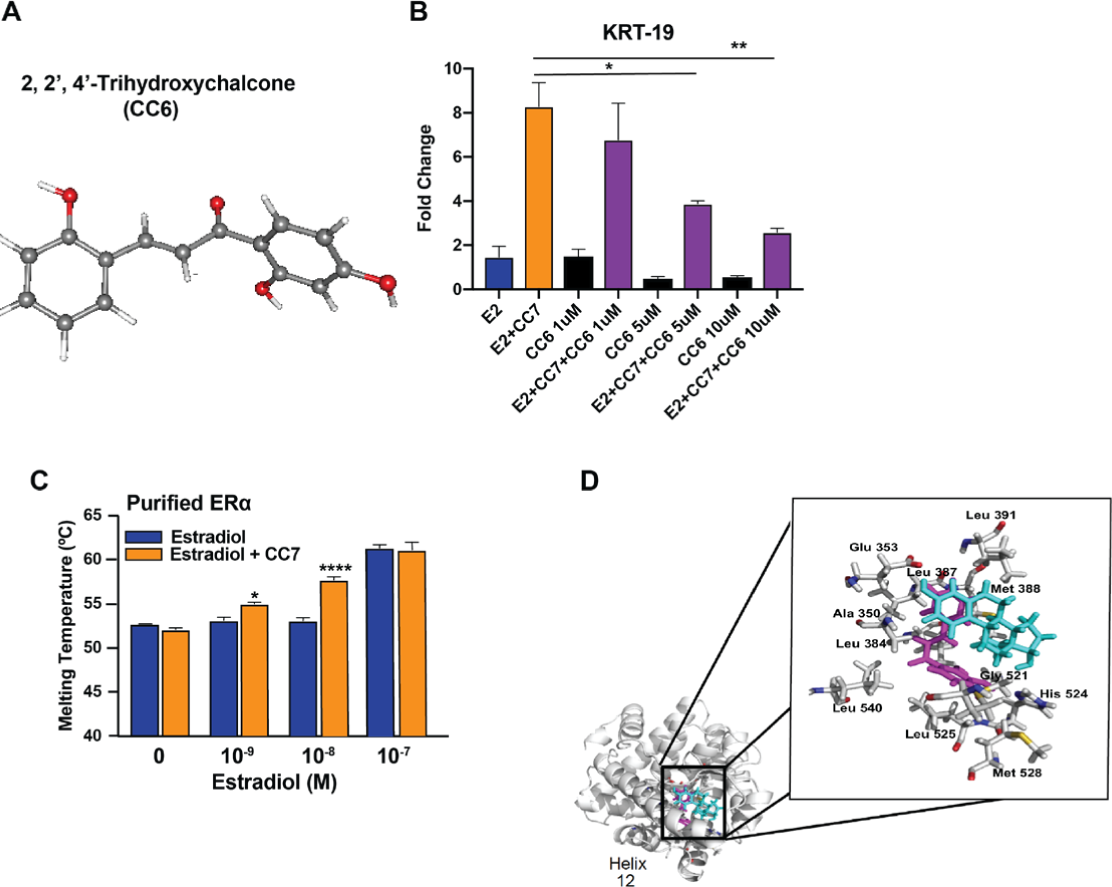
CC7 binds in the binding pocket of ERα in close proximity to E2 by molecular simulation studies. **(A)** Chemical structure of CC6 (adapted from PubChem). **(B)**KRT-19 gene expression in U2OS-ERα cells determined by RT-qPCR after cells were treated with E2 (10 nM), CC7 (5 μM) alone or in combination with increasing doses of CC6 for 24 hrs in technical triplicates (**C**) Thermostability assays performed with purified ERα LBD in increasing concentrations of E2 without or with 5 μM of CC7. The significant differences between various amounts of E2 alone and in combination with CC7 were analyzed with Two-way ANOVA followed by Sidak’s multiple comparisons post hoc test. (**D**) Molecular dynamics simulations of predicted interactions for CC7 in adducts with the crystal structure of the ligand binding domain of estrogen receptor complexed with ligand 17 beta-estradiol (PDB accession number: 1ERE) at 12ns. ERα and the corresponding CC7 were subjected to MD simulations using Desmond and visualized using the molecular graphics program PyMol. Estradiol is depicted in cyan, CC7 is depicted in magenta, and close contact residues within 3 Å of 2’, 3’, 4’-THC are shown in white.

To further evaluate our coligand model, we performed thermostability assays to measure the melting temperature of purified ERα LBD (Fig. 3C). In this assay, changes in melting temperature only occur when the ligand alters conformation. E2 at 1 and 10 nM did not produce a significant change in melting temperature, whereas an increase was observed at 100 nM. CC7 had no effect on the melting temperature of ERα LBD alone, but produced an increase in the presence of 1 and 10 nM E2. Using coordinates from the crystal structure of E2 complexed to the ERα LBD^16^ we simulated the binding of both CC7 and E2 as coligands (Fig. 3D). Modeling with E2 positioned in its binding site, CC7 interacts with a secondary site within the ligand pocket using contact points located in helix 3 (Met 343, Ala 350, Glu 353), helix 6 (Leu 384, Leu 387, Leu, 391), helix 11 (Gly 521, Leu 525, His 524, Met 528, and helix 12 (Leu 540) (Fig. 3D, inset). Computer simulations predict that coordinate binding of CC7 and E2 produces a different conformation of the ERα LBD than with E2 alone (Supplementary Fig. S2 and S3). The most dramatic change being a predicted shift in residues 360-370, 460-475, and the N-terminus (Supplementary Fig. S2 and S3).

### CC7 acts as an ER antagonist in MCF-7 breast cancer cells

Current MHT treatments lead to increases in cancer risk. Thus, we next assessed the effects of CC7 alone and in combination using MCF-7 breast cancer cells. Expectedly, E2 increased luciferase reporter activity on an ERE whereas CC7 remained inert, showing no increase in luciferase activity. Interestingly the combination of E2 with CC7 did not synergize luciferase activity in MCF-7 cells (Fig 4A). Furthermore, CC7 decreased proliferation of MCF-7 cells in a dose-dependent manner (Fig 4B). Treatment of MCF-7 cells with E2 expectedly increased MCF-7 cell proliferation whereas CC7 alone had no effect on proliferation (Fig. 4C). CC7 blocked proliferation induced by E2 in a dose-dependent manner (Fig. 4C). In addition to reducing cell number, CC7 blocked progression of cells into S-phase, as measured by flow cytometry (Fig. 4D). The antiproliferative action of CC7 is likely mediated by ERα, because these MCF-7 cells do not express ERβ^9^.

**Figure 4.**
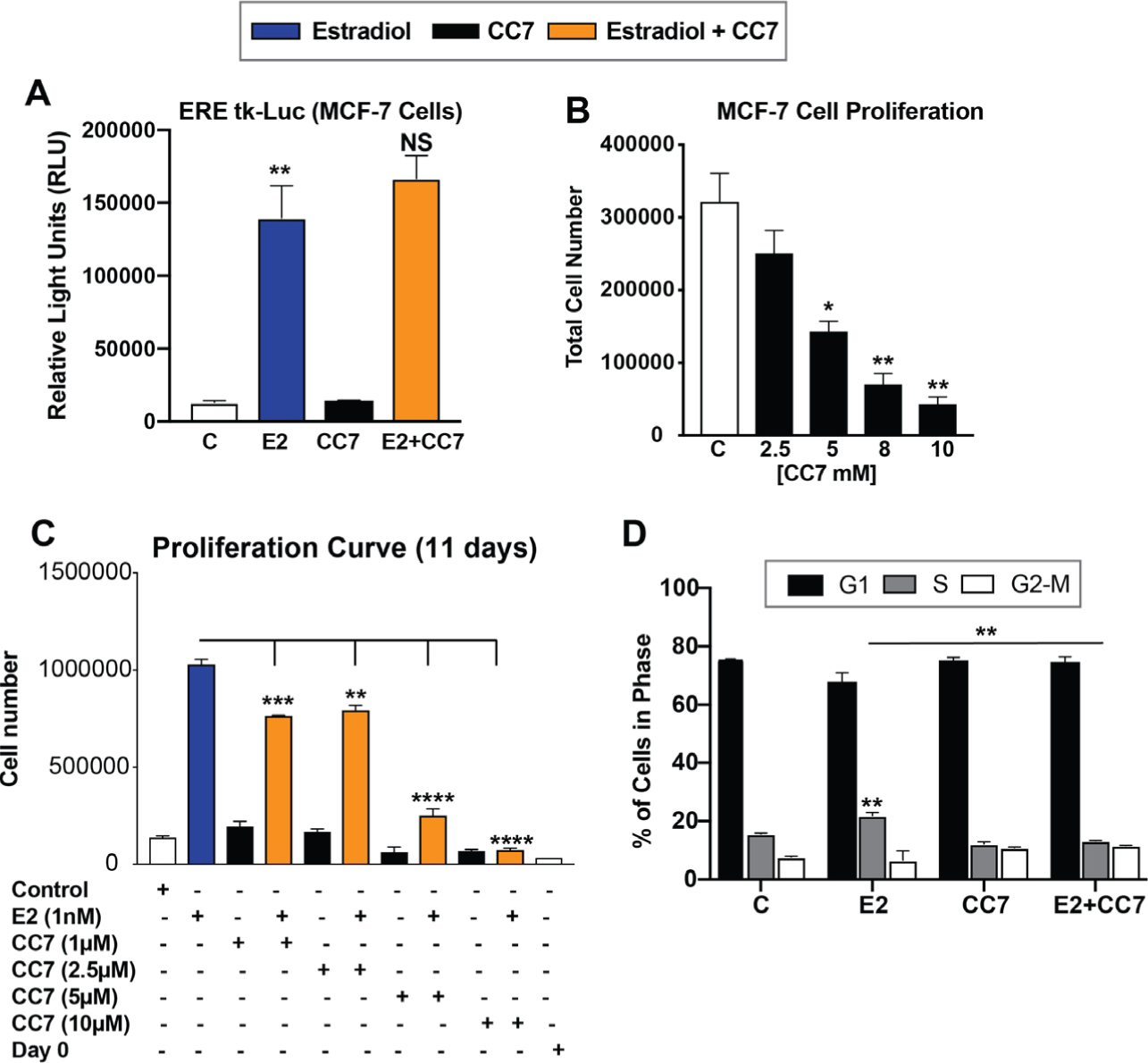
E2 induced MCF-7 breast cancer cell proliferation was blocked by CC7. (**A**) Measured RLU in MCF-7 cells transfected with an ERE-tk Luciferase and treated for 24hrs with 10nM estradiol, 5 μM CC7 or CC7/E2 combination. (**B**) Cell numbers counted after treatment with increasing concentrations of CC7 and **(C)** Cell numbers counted after treatment with increasing doses of CC7 without or with E2 (10 nM) for 11 days. (**D**) % of cells in G1, S or G2-M cell cycle phases. Cells determined by Flow Cytometry analysis after treatment with vehicle control, E2, CC7 or the combination for 24 hours. Error bars are means ± SEM.

### CC7 acts as an ERα antagonist in the mouse uterus

To explore the effects of CC7 in vivo on estrogen-sensitive tissues, 8 wk old C57BL/6 female mice were ovariectomized or left intact and then were treated with estradiol, CC7 or the combination for 4 weeks using an osmotic pump implanted posterior to the scapula (for OVX mice) or just CC7 (for intact mice) by IP inectionfor 4 weeks and the size of the uterus was measured. In intact mice, CC7 decreased uterine weight (Fig 4A).In OVX mice, CC7 did not increase uterine size, whereas E2 produced a large increase in uterine weight (Fig. 5B). CC7 blocked the E2-induced increase in uterine weight (Fig. 5B). CC7 blocked known estrogen induced transcripts in the uterus, S100A9 and LCN2^17^ (Fig. 5C). Hematoxylin and eosin staining showed that compared to the control mice CC7 did not increase the number of cells lining the ducts (Fig. 5D). E2 increased the layer of cells lining the duct and CC7 inhibited the proliferative response (Fig. 5D). In contrast, CC7 failed to antagonize the E2-mediated decrease in gonadal adipose (Fig. 5E). We observed a significant decrease in in body weight in animals treated with estradiol alone, CC7 alone or the combination compared to vehicle treated animals. (Fig. 5E). In contrast, no changed in gonadal adipose tissue weight or mammary weight were observed (Fig 5E).

**Figure 5.**
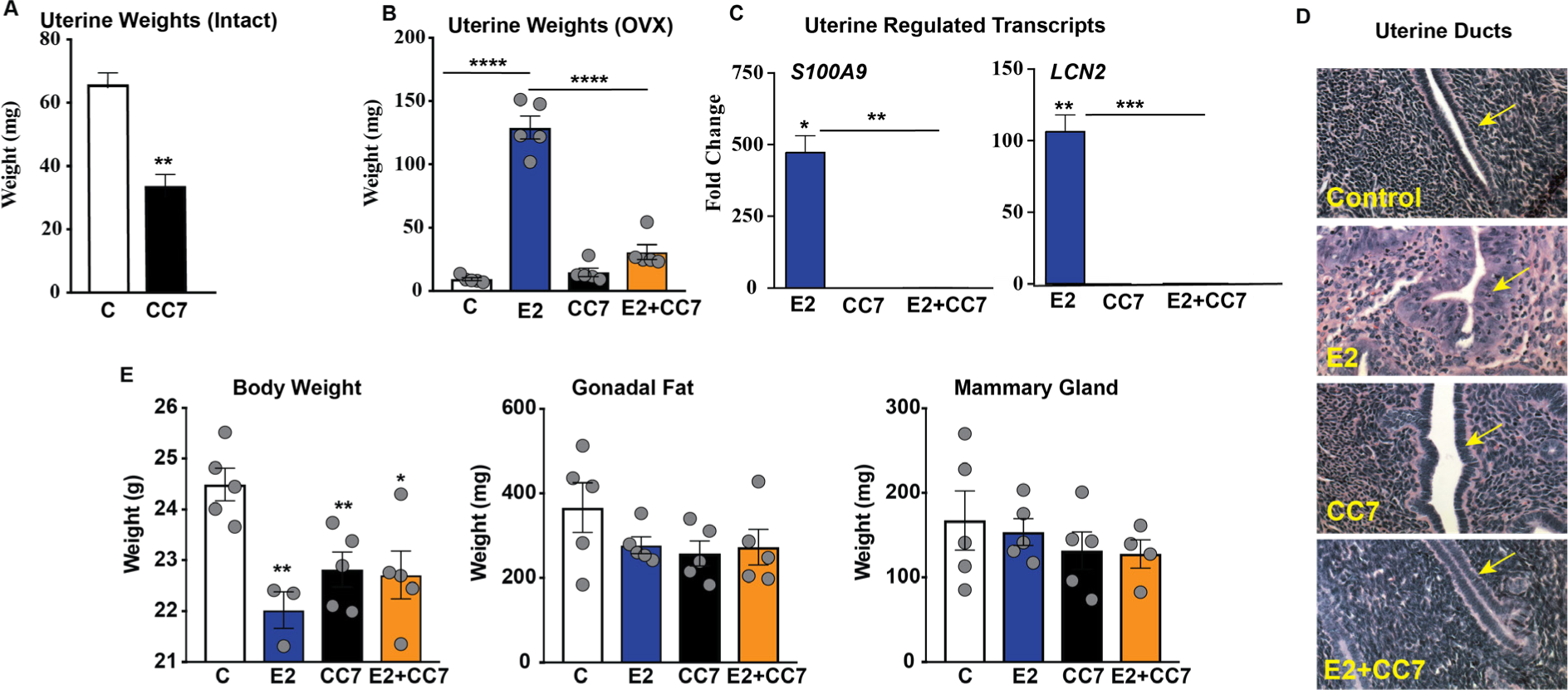
CC7 acts as an ERα antagonist in the mouse uterus. (**A**) Uterine weights of intact female mice that recieved vehicle control or 2mg CC7 for 3 wks by IP injection (**B**) Uterine weights of OVX mice received E2, CC7 alone or in combination using an osmotic pump for 4 weeks. The asterisks indicate the significant difference between E2 alone and the combination determined with One-way ANOVA followed by Tukey’s multiple comparisons post hoc test. (**C**) E2 regulated transcrip levels in uterine tissue of OVX mice treated with E2 alone or in combination with CC7 for 4 weeks determined by RT-qPCR. (**D**) H&E staining of sectioned uterine tissue. (**E**) Body weights, gonadal adipose tissue weights and mammary gland weights of OVX mice treated with E2, CC7 alone or in combination for 4 weeks. Error bars are mean ± SEM. Data was analyzed with one-way ANOVA followed by Turkeys multiple comaprisons post hoc.

## DISCUSSION

In postmenopausal women, MHT effectively prevents osteoporosis and bone fractures^18^, and reduces body weight^19^, obesity^20^ and type 2 diabetes^21,22^. Discovering strategies that render MHT safe for continuous, long-term therapy by overcoming the proliferative actions of ERα, poses a pharmacological challenge. In our approach, we screened for compounds that could bind simultaneously with E2 to ERα, but at a coligand site on the receptor, to create a two-ligand, one-receptor mechanism. Our screen identified CC7 as a potential ERα coligand that reprograms E2 when bound to ERα. The reprogramming of E2 could result in optimized transcriptional regulation, cellular activity and tissue modulation to preserve and enhance desirable, while minimizing its undesirable effects.

E2 produces tissue specific effects through a variety of mechanisms. In different tissues or cell types like U2O2 and MCF-7, ERα binding elements differ in their availability^23^. Additionally, tissues express different transcription factors. For instance, FoxA1 is involved in E2-mediated transcription of ERα target genes in MCF-7 cells, but U2OS-ERα cells lack FoxA1 expression^23^. Furthermore, transcriptional coregulators express differently among different tissues^1^. Not surprisingly, cotreatment of CC7 and E2 causes a cell and tissue-specific reprogramming of E2 bound ERα. In U2OS-ERα cells, CC7 potentiates the effects of E2 in transfection assays and produces a synergistic activation of known E2 target genes, KRT19 and NKG2E, but was relatively inactive alone. Cotreatment with CC7 and E2 reprogrammed E2 gene regulation, affecting over 800 unique genes in U2OS-ERα cells. In U2OS cells, CC7 does not exhibit E2 antagonistic activity like the SERM. Instead, CC7 functions as an ERα coagonist by changing the magnitude and type of genes regulated by E2. In a different model, MCF-7 cells, CC7 blocked the agonistic effect of E2 on cell proliferation and gene expression. As our MCF-7 cells do not express ERβ, the effects of CC7 must be mediated by ERα, Like MCF-7 cells, CC7 also exhibited an antagonistic action on the uterus in ovariectomized mice treated with exogenous E2. However, CC7 had no antagonistic effect on the E2-induced reduction of gonadal adipose tissue, suggesting either a coagonist or third type of effect in adipose tissue. Much like the effects of E2 alone, our observations suggest that differential tissue expression of transcription factors, such as FoxA1 or coregulatory proteins could account for the tissue-specific regulation by the CC7/E2 combination.

E2 binding to the LBD produces a conformational change that creates a binding surface for coregulators^25^. The observation that the size of the binding cavity is nearly two times larger than the molecular volume for E2^16^, suggests that two ligands, such as E2 and CC7 can occupy the pocket at the same time. Our studies suggest that E2-induced conformational changes create a coligand binding site (CBS). Once E2 is bound the modeling shows that CC7 binds to a single site in close proximity to E2. The presence of two ligands in the pocket creates a conformation distinct from E2 bound ERα, which could alter the regulatory elements it binds to and recruitment of coregulators.

Although it is possible that E2 and CC7 form a heteroligand, as described with cotreatment with E2 and SERMs^24^ our data from several studies is consistent with our coligand model. First, thermostability studies using pure ERα LBD showed that E2 produced an increase in melting temperature at 1 and 10 nM only when CC7 was present. Since the reaction only contained the pure ERα LBD and the ligands, the only feasible way that CC7 could change the melting temperature of the E2-ERα LBD complex would be by binding to the LBD along with E2. Next, in competition studies, we found that low concentrations of CC7 increased [3H]-E2 binding. One explanation for this finding is that CC7 binding to the CBS interferes with the exit of [3H]-E2 from the binding cavity. Increased E2 binding could also account for the enhanced recruitment of ERα by ChIP in U2OS cells. At 100 μM, CC7 started to compete with [3H]-E2 binding, suggesting that at very high concentrations, CC7 can bind to the E2 binding site, preventing [3H]-E2 binding. In addition, the observation that another structurally related chalcone, CC6 blocked the synergistic action on gene expression without blocking the effects of E2, suggests that it blocks only CC7 binding to the CBS and not the binding of E2 to its site. Finally, our studies showed that in the absence of E2, CC7 is inactive, probably because it cannot bind to ERα since no CBS exists. Thus, multiple lines of experimental evidence support our one receptor, two ligand model, which is consistent with the modeling data. Future X-ray crystallography studies will be needed to prove the one receptor, two ligand model and determine the precise location of the CBS for CC7. Such studies could be very challenging, because crystallization will require both CC7 and E2 to be bound simultaneously to ERα.

Both CC7 and tamoxifen antagonize the effects of E2 on MCF-7 cell proliferation, however their mechanisms are different. SERMs produce their antagonist effects by competitively inhibiting E2 binding to ERα and recruiting corepressors when ERα is bound to DNA^26^. In contrast, CC7 actually increased [3H]-E2 binding to its pocket where it remains operative, but results in reprogrammed gene regulation.

Chalcones are a structurally diverse group of flavonoids synthesized by many plants. A potential important ramification of our study could be that chalcones, and possibly other plant compounds commonly ingested in human diets, could act as coligands to alter developmental effects mediated by ERα during female growth and maturation, as well as diseases associated with the natural changes of estrogen concentrations during the menstrual cycle and the decline in circulating estrogens during menopause. The presence of coligands in different geographic diets could be involved in the observation that the incidence of breast cancer varies greatly in different counties^27^.

CC7 exhibits several pharmacological properties potentially favorable for long-term MHT, in postmenopausal women. Because it antagonizes the proliferative^13^ effects of E2 on MCF-7 breast cancer cells and the mouse uterus, CC7 could possibly be used in combination with existing estrogens in MHT, instead of progestins, to prevent the increased risk of cancer associated with MHT. It is also conceivable that the combination of CC7 and E2 result in lower E2 dose required to exert its pharmacological effect, closer to known pre-menopausal circulating serum E2 levels. Our results showing that CC7 potentiates E2 transcriptional activation of ERβ might be a useful property to further prevent the proliferative effects of estrogens. We previously showed that the ERβ-selective plant extract^28^, MF101, reduced hot flashes in postmenopausal women,^29,30^ suggesting another possible indication for CC7. Our findings suggest that an ERα coligand/estrogen combination could potentially provide superior safety to the estrogen only and estrogen/progestin regimens, and might make the current estrogens in MHT safe for long-term therapy to prevent chronic diseases associated with menopause.

Different ligands that bind to the same binding pocket of ER and other nuclear receptors create different conformations leading to the recruitment of distinct coregulators to alter gene expression and biological effects^31–33^. A major pharmaceutical strategy to overcome side-effects is to design and synthesize selective nuclear receptor modulators that bind to the same binding pocket as the cognate ligand, but produce different conformations and biological responses^34,35^. The use of coligands that bind to a secondary site in the ERα LBD, and potentially other nuclear receptors, offers a new strategy to improve drug safety by reprogramming the actions of an endogenous steroid bound to its nuclear receptor. Generalization of coligands to other nuclear hormone receptors, each with slightly different LBD, and therefore CBS, would open a multitude of possible diet/hormone interactions as well as therapeutic possibilities.

## MATERIALS & METHODS

### Compounds

2’, 3’, 4’-trihydroxychalcone and 2, 2’, 4’-trihydroxychalcone were obtained from INDOFINE Chemical Company (Hillsborough, NJ). All other compounds were obtained from Sigma Aldrich (St. Louis, MO). The chemicals were dissolved in ethanol and used at a final concentration of 0.1%.

### Cell Culture

Human U2OS cells expressing a tetracycline-regulated ERα (U2OS-ERα) and ERβ (U2OS-ERβ) were prepared, characterized, and maintained as previously described36. The cells were maintained in phenol red-free DMEM/F-12 (Gibco, Life Technologies) supplemented with 5% charcoal-dextran stripped fetal bovine serum (Gemini Bio-Products, Sacramento, CA), 100 units/mL penicillin and streptomycin, 50 μg/mL fungizone and 2 mM of glutamine. To maintain stable transfected cells, 50 μg/mL hygromycin B (Invitrogen) and 500 μg/mL of zeocin (Invitrogen) were included in culture media. MCF-7 breast cancer cell line obtained from ATCC was maintained in phenol redfree DMEM/F-12 supplemented with 10% fetal bovine serum (Gemini Bio-Products), 100 units/mL of penicillin and streptomycin, 50 μg/mL fungizone and 2 mM of glutamine. For experiments, the culture media were replaced with 5% stripped FBSDMEM/F12.

### Transient Transfection Assays

U2OS cells (wild type) were transfected with 3 μg of a plasmid containing the ERE upstream of the minimal thymidine kinase luciferase promoter and 1 μg of CMV-ERα or CMV-ERβ by electroporation as previously described37. The transfected cells were treated with various reagents for 24 hours and then lysed and assayed for luciferase activity using the Luciferase Assay System (Promega Corp., Madison, WI) according to the manufacturer’s protocol.

### Competitive Estrogen Receptor Binding Assays

U2OS-ERα cells grown in 12-well dishes were treated with or without doxycycline (1 μg/ml) for 24 hours. The cells were then incubated with 5 nM [3H]-E2 (specific activity 87.6 Ci/mmol; PerkinElmer Life Science, Waltham, MA) in the presence of increasing concentrations of 2’, 3’, 4’-THC at 37°C for 1 hour as previous described36. After washing the cells with 0.1% bovine serum albumin in PBS, 100% ethanol was added. Radioactivity was measure in the samples with a scintillation counter. Specific binding of [3H]-E2 was calculated as the difference between total and nonspecific binding in counts per minute (CPM).

### Cell and Tissue Total RNA Extraction and RT qPCR

Total cellular RNA was extracted using the Aurum Total RNA Mini Kit (Bio-Rad Laboratories, Hercules, CA) following the manufacturer’s protocol. Animal tissues were dissected and immediately frozen in liquid nitrogen. To isolate RNA, the frozen tissues were first homogenized in PureZOL RNA isolation reagent (Bio-Rad) using MP FastPrep-24 (MP Biomedicals, Santa Ana, CA). Total RNA was then extracted using the Aurum Total RNA Mini Kit for Fatty and Fibrous Tissue (Bio-Rad). Reverse transcription reactions were performed using the iScript cDNA Synthesis Kit (Bio-Rad) with 1 μg of total RNA according to the manufacturer’s protocol. Quantitative PCR was performed with Bio-Rad CFX96 Thermal Cycler System using SsoFast EvaGreen Supermix (Bio-Rad). The results were analyzed by competitive Ct method38. The Ct values of specific genes were adjusted by GAPDH (for human cell lines) or Gus (for mice tissues) running simultaneously to obtain ΔCt. The fold changes were computed by comparison of adjusted Ct values (ΔCt) from various treatments with control samples. All data presented are mean ± SEM.

### Microarray and Data Analysis

Total cellular RNA was isolated utilizing the Aurum RNA isolation kit (Bio-Rad, Hercules, CA) per the manufacturer’s directions. RNA isolates were first quantified by nanodrop, and then qualitatively evaluated by the Bio-Rad Experion system per the manufacturer’s instructions. Biotin-labeled cRNA samples were prepared using 750 ng of total RNA. Biotin-labeled samples were evaluated by both 260/280 absorbance spectrophotometry and capillary electrophoresis. Labeled cRNA samples were hybridized overnight against Human genome HG U133A-2.0 GeneChip arrays, (Affymetrix, Santa Clara, CA). All treatments were done in triplicate with the same batch of microarrays. The data was analyzed as previously described17.

### Chromatin Immunoprecipitation (ChIP)

ChIP assay was performed as previously described^39^. U2OS-ERα cells were incubated for 24 hours with 1 μg/ml of doxycycline in serum-free DMEM/F12 upon 80% confluence. The cells were then treated with vehicle, E2 (10 nM), 2’, 3’, 4’-THC (5 μM) and the combination for 1 or 2 hours. For MCF-7 cells, the cultured media were switched to serumfree DMEM/F12 upon 80% confluence and incubated for 24 hours. The cells were then treated with E2, 2’, 3’, 4’-THC and in combination for 1 or 2 hours. To end treatments, the cells were fixed by adding 11X formaldehyde solution to the culture media and incubated for 15 minutes at room temperature with shaking. The reaction was quenched with 1.25M glycine solution. The cell monolayer was then washed with PBS containing cOmplete Protease Inhibitor Cocktail (Roche), collected by scraping and concentrated by centrifugation (2000xg, 4°C for 5 minutes). The cell pellets were saved at -80°C. To perform the ChIP assay, the frozen pellets were thawed on ice and lysed with buffer containing 0.5% of Triton X-100, 50 mM of Tris (pH 7.4), 150 mM of NaCl, 10 mM of EDTA and Protease Inhibitors. Cell lysates were centrifuged and pellets were resuspended in RIPA buffer. The suspensions were sonicated on ice using a Digital Sonifier (Brason) and supernatants were obtained by centrifuge at 14000 rpm for 10 min at 4°C. The samples were then diluted with appropriate amount of dilution buffer (RIPA buffer without detergents). Approximately 10% of each diluted sample was taken for input and stored at 4°C. The samples were incubated with 4 μg/ml of rabbit anti-ERα IgG (sc-544, Santa Cruz Biotechnology) at 4°C overnight with rotation. The immune complexes were then precipitated by Protein G Magnetic Sepharose beads (GE Healthcare) for 4 hours rotating at 4°C. The DNA-protein-antibody complexes were then eluted from magnetic beads with elution solution (1% SDS, 0.1M NaHCO3) at 65°C for 10 minutes. The bound DNA was reverse cross-linked by incubation at 65°C overnight. Eluted DNA was purified and concentrated using the ChIP DNA Clean and Concentrator (Zymo Research, Irvine, CA). ERα antibody precipitated DNAs were amplified by real-time PCR with specific primers for the ERE in KRT19^15^. The Ct values from treatments were adjusted by corresponding input Ct values. The fold changes were obtained by comparison of adjusted Ct values of treatments with that of control. The results are expressed as means ± SEM.

### Cell Proliferation Studies

Cells were plated at a density of 50,000 cells per well in 6-well tissue culture plates in DMEM/F12 supplemented with 5% stripped FBS. At following day, the cells were treated with vehicle, E2 (1 nM) in the absence and presence of increasing doses of 2’, 3’, 4’-THC (1, 5 and 10 μM) for 5 days. The cells were then detached with trypsin, neutralized with media containing 5% FBS and well suspended by pipetting and vortex. The cell numbers were then counted using a Coulter Counter (Particle Counter Z1, Beckman). All treatments were biological triplicates and presented as mean ± SEM.

### Flow Cytometry Analysis

Flow Cytometry was carried out base on previously described method39. Briefly, the cells were plated at a density of 500,000 cells per well in 6-well tissue culture dishes in DMEM/F-12 supplemented with 5% stripped FBS for 48 hours. The cultured medium was then replaced by serum-free DMEM/F12 for 24 hours. The cells were then treated with vehicle, E2 (0.1 nM) without or with indicated amount of 2’, 3’, 4’-THC for 24 h. The media were then aspirated and the cell monolayer was washed with PBS, detached with trypsin and collected by centrifugation at 1700 rpm for 5 minutes. The cell pellets were washed with ice cold PBS followed by centrifuge at 1700 rpm for 10 minutes at room temperature. The cell pellets were resuspended in 500 μl PBS containing 50 μg/ml propidium iodide, 0.1% of triton X-100, 0.1% of sodium citrate, and 10 μg/ml of RNase. The cell suspensions were then run on BD LSR II Flow Cytometer (BD Biosciences) in the Flow Cytometry facility at UC Berkeley and cell cycles were analyzed using FlowJo 7.6.5.

### Molecular Dynamics Simulation

The structure of the ERα LBD domain was obtained from the Protein Data Bank with accession number 1ERE. The PRODRG server was used to produce the structure files for modeling the THC structure. The ERα LBD was processed in Protein Preparation Wizard (Schrödinger Suite 2015-4 Epik version 3.4, Schrödinger, LLC, New York, NY, 2015; Impact version 6.9, Schrödinger, LLC, New York, NY, 2015; Prime version 4.2, Schrödinger, LLC, New York, NY, 2015) to add hydrogens, assign bond order and other basic PDB repair tasks. The crystal structure of ER□ LBD under Prime version 4.2 (Schrödinger, LLC, New York, NY, 2015), to repair missing sidechains, gaps and loops (His 377, His 501, Gln 506, His 513, and Cys 530). The structure was then treated with a molecular dynamics (MD) simulation of 12 ns using DESMOND Molecular Dynamics System, version 4.4 (D. E. Shaw Research, New York, NY, 2015. Maestro-Desmond Interoperability Tools, version 4.4, Schrödinger, New York, NY, 2015) so as to explore the structure landscape. In the Desmond simulation we used the OPLS all atom force field. The protein structure was solvated with a simulated SPC water model in an orthorhombic box with buffer space of 10 Å from the edges of the protein. The system was equilibrated by incorporating the default Desmond Relax protocol in Maestro. The simulation ran for 12ns using Desmond at 300K/1atm in the NPT ensemble employing a Nose-Hoover thermostat and a Martyna-Tobias-Klein barostat with a 2.0ps relaxation time. Coulomb interactions were assessed using a 9 Å short range cutoff and smooth particle mesh Ewald as a long-range method where Ewald tolerance= 10-9. MD simulation was analyzed using the analytical tools in the Desmond package. The potential energy of the protein as well as the total energy of the entire system was calculated using the MD simulation quality analysis. The stability of the docked complex across the trajectory was assessed from their RMSD (root mean square deviation) plots. The RMSF plot (root mean square fluctuation) of the backbone and C-α atoms of each residue from its time-averaged position was also generated.

### Protein Thermostability

Purified ERα LBD containing amino acids 315-545 used by Brzozowski et al for crystallization16 with a his-tag was custom made (GenScript). All reactions were set up in 20 μL reactions in 96 well plates with purified ERα LBD at a concentration of 0.6 μg/μL and 10x SPYRO orange dye (Invitrogen). The ERα LBD was incubated with a concentration of 5 μM 2’, 3’, 4’-THC and increasing doses of E2 (1 nM, 10 nM, 100 nM) and compared to E2 doses alone. Thermal melting experiments were performed on the Viia7 instrument (Life Technologies) melt curve program with a ramp rate of 0.05°C/s, and the ramp goes to 95°C. Melting temperatures were analyzed with Protein Thermal Shift Software (Life Technologies) to identify the midpoint of the transition with an analysis of the first derivative. All experiments were performed with triplicate samples.

### Animal Studies

Eight week old C57bl/6J female ovariectomized or intact mice were purchased from Jackson Laboratories (Sacramento, CA). Mice were housed and maintained according to the Office of Laboratory Animal Care (OLAC) standard procedures in the Northwest Animal Facility at UC Berkeley. All mice were fed a soy-free chow diet (Harlan Laboratories, Livermore, CA) starting one week before IP Inection or osmotic pump implantation. For IP inection mice were injected daily for 3 weeks with vehicle control or 2mg of 2’, 3’, 4’-THC. Mini-Osmotic Pumps (model 2006) were purchased from Alzet (DURECT Corporation, Cupertino, CA). The pumps were filled with vehicle, 1 μg of 17β-estradiol, 2 mg of 2’, 3’, 4’-THC or the combination of 2’, 3’, 4’-THC with E2. All drugs were dissolved in sterile vehicle consisting of 50% DMSO, 25% ethanol and 25% deionized water. Pumps were handled with sterile gloves and filled using a 27 gauge filling tube and 1 mL syringe. All pumps were placed in 1X PBS in 15 mL sterile conical tubes and incubated overnight at 37°C. Pumps were surgically implanted into 8 week old C57bl/6J female ovariectomized mice posterior to the scapula and left for a duration of 4 weeks. Intact mice were injected intraperitoneal with 2’, 3’, 4’-THC every day for 3 weeks. Mice were weighed once a week for the duration of the experiment. The intraperitoneal gonadal fat was removed and weighed. Uterine tissue was collected, fluid drained and then weighed. For uterine histology, tissue was removed and trimmed of excess adipose tissue. Tissues were fixed in formalin for 24 hours then transferred to 50% ethanol for 1 hour followed by 70% ethanol for another hour. After fixation tissues were sent to Histopathology Reference Laboratory (Hercules, CA) where they were paraffin embedded, sectioned and stained with hematoxylin and eosin for morphological examination.

### Statistical Analysis

All data are presented as the mean ± SEM from at least biological triplicates. The statistical significance of the difference between two groups was assessed by Student’s t-test. For the data sets consisting of more than two groups the statistical significance of differences among various groups (treatments) were analyzed by one-way analysis of variance (one-way ANOVA) tests or two-way ANOVA as specified in figure legends. All ANOVA tests were followed by Tukey’s or Sidak’s multiple comparisons post hoc tests to analyze the significance of differences between any two different groups (treatments) or control as indicated in figure ledged. Statistical analysis and graph plotting were performed using GraphPad Prism version 6 (GraphPad Software Inc.; La Jolla, CA, USA). The statistical significance for the numbers of asterisks in the figures are *p<0.05; ** p<0.01,*** p<0.001, and **** p<0.0001.

