## Supplementary figures and images for "2’, 3’, 4’-trihydroxychalcone is an Estrogen Receptor Ligand Which Modulates the Activity of 17β-estradiol"

### Supplemental Figure 1

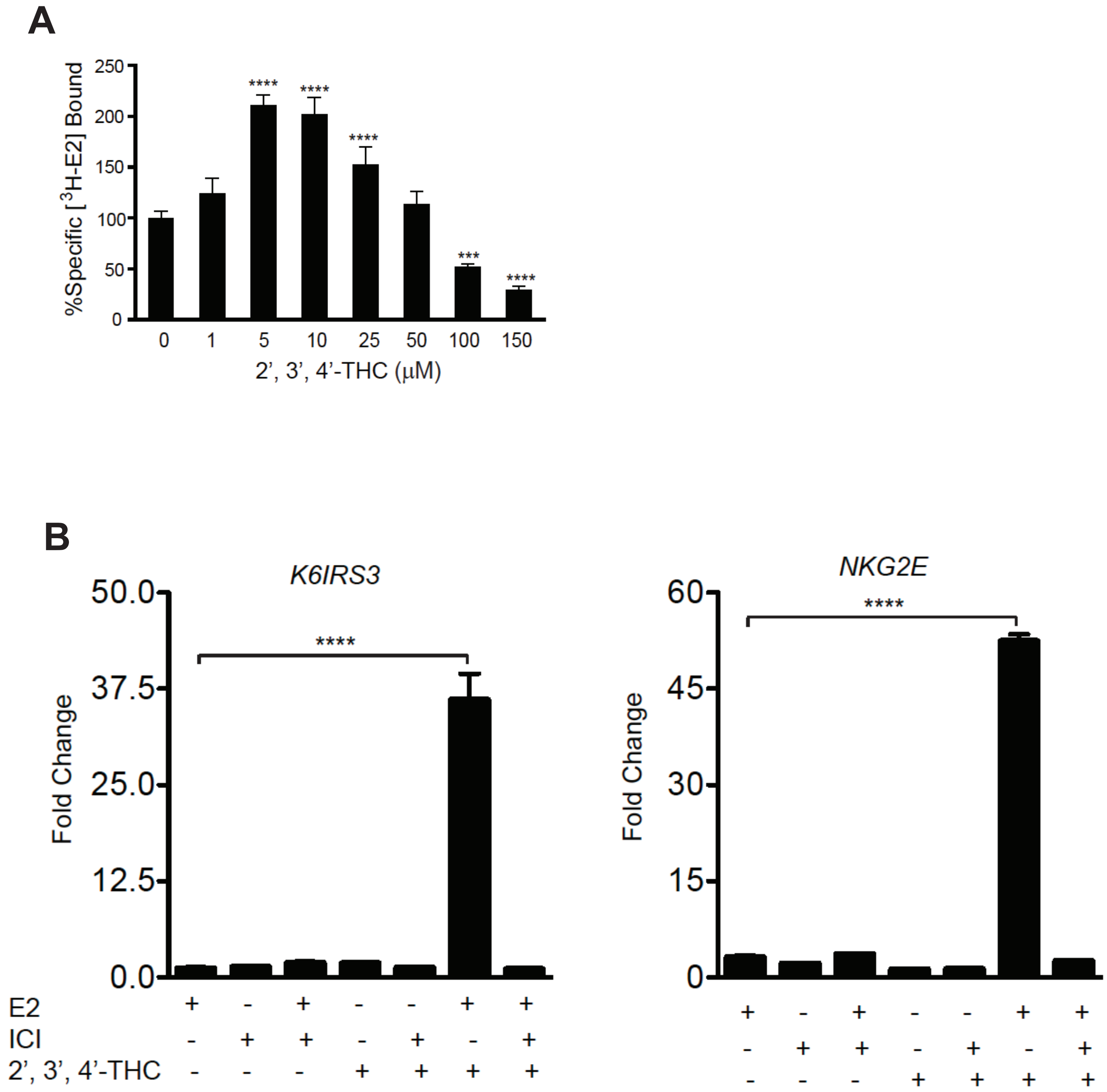


**Supplemental Figure 1**

### Supplemental Figure 2

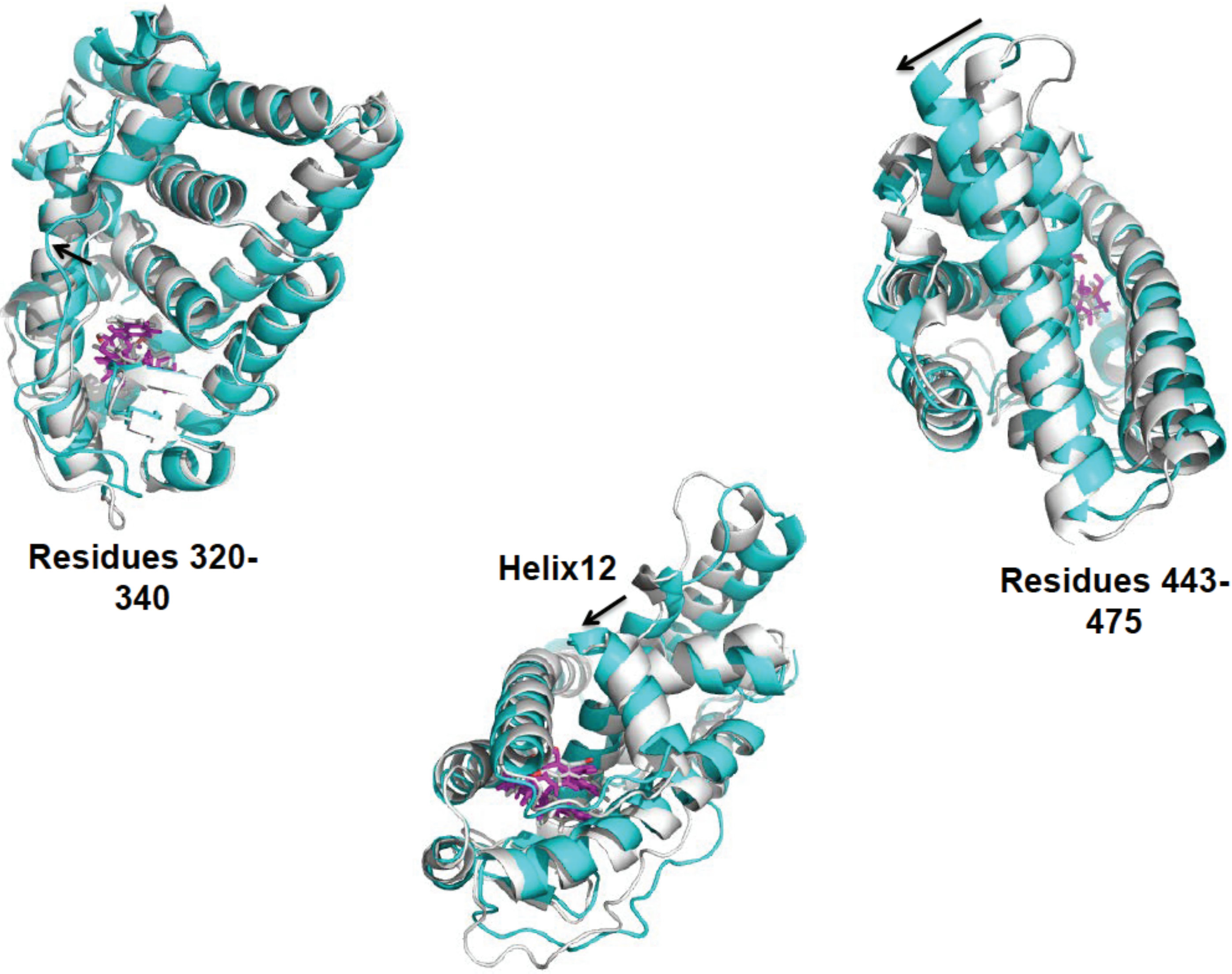


**Supplemental Figure 2**

### Supplemental Figure 3

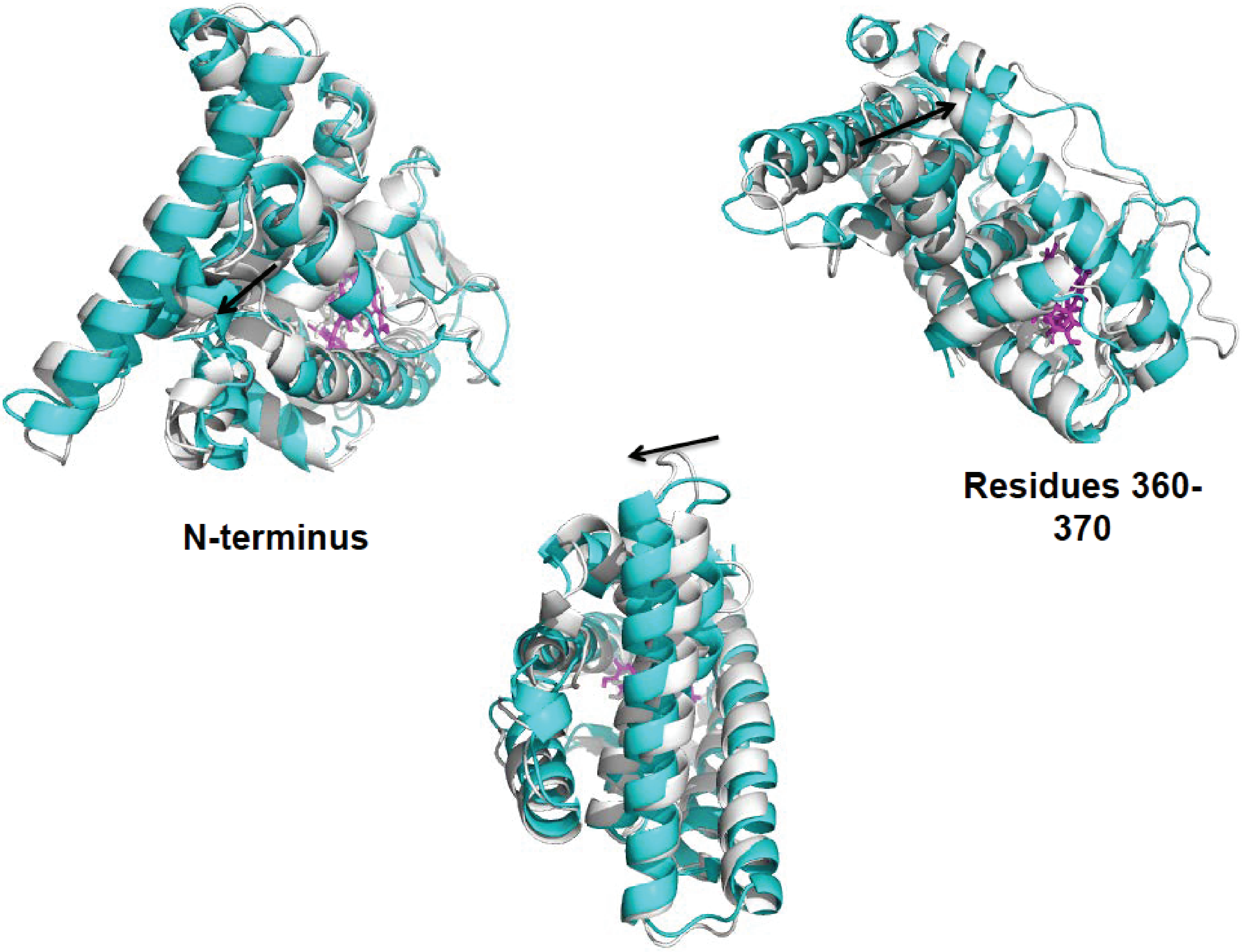


Supplemental Figure 3
