## Supplemental Table 1 for "2’, 3’, 4’-trihydroxychalcone is an Estrogen Receptor Ligand Which Modulates the Activity of 17β-estradiol"

| | **Class 1:** 2', 3', 4'-THC Antagonizes | **Class II:** 2', 3', 4'-THC Potentiates | **Class III:** Newly Regulated Genes |
|---|---|---|---|
| **Up-regulated by estradiol** | 24 | 75 | |
| **Down-regulated by estradiol** | 5 | 20 | |
| **Activates** | | | **327** |
| **Represses** | | | **268** |

### Supplementary Table 1
